# Rethinking synthesis and function of neuro-estrogen and neuro-androgen in the hippocampus: some methodological problems and possible solutions

**DOI:** 10.1101/2025.08.13.670028

**Authors:** Suguru Kawato, Mari Ogiue-Ikeda, Mika Soma, Shigeo Horie, Minoru Saito, Kouichi Minato

## Abstract

Local synthesis and action of neuro-estrogen and neuro-androgen (including neuro-estradiol (nE2) and neuro-testosterone (nT) have become recognized as key mechanisms in modulation of neural plasticity and cognitive performance, in addition to the contribution of circulating sex steroids. However, still several methodological problems are left to be solved in order to get an better understanding of functions of neural sex steroids in the brain. Here we describe and discuss important improvements in the methods for determination of accurate concentrations of nE2 and nT in rat hippocampus (Section 1), and in methods for analysis of the dendritic spine density in castrated male rat hippocampus (Section 2).

The improvements are discussed in order to solve the following two problems. One problem is that the previously reported nE2 concentrations in the hippocampus were widely distributed between ∼3 pg/g tissue and ∼2 ng/g tissue (i.e., between ∼11 pM and ∼7 nM) (Section 1). The other problem is that the degree of decrease in hippocampal spine density by castration in male rat hippocamps was strongly dependent on surgical skills of operators (surgeons) (Section 2).

## Section 1: Rethinking concentration of nE2 and nT in hippocampus

### Introduction 1

Local synthesis of nE2 has been attracting much attention and extensively studied in the hippocampus, center for learning and memory. However, the determination methods for accurate concentration of hippocampal nE2 still have difficulties. The reported nE2 concentrations in recent 20 years show significant differences between different laboratories, for example, even concerning only the hippocampus of adult rats and mice. Interestingly, previously reported nE2 concentrations in the hippocampus of male rodents were widely distributed between ∼3 pg/g tissue and ∼2 ng/g tissue (between ∼10 pM and ∼8 nM) depending on different publications (Hojo et al 2009) (Caruso et al 2013) (Li & Gibbs 2019). It should be noted that, on the other hand, previously reported nT concentrations in male hippocampus were around ∼ 3 ng/g tissue (∼10 nM) which are not so much different between previous reports (Hojo et al 2009) (Caruso et al 2013) (Li & Gibbs 2019).

In this Section 1, we measured nE2 and nT concentrations in frozen-thawed hippocampal tissues, in order to compare them with nE2 and nT concentrations in freshly prepared (not frozen-thawed) hippocampal tissues. We observed a considerable difference in nE2 concentration in frozen-thawed hippocampal tissues compared to fresh hippocampal tissues. However, nT concentration was not considerably different between these two types of hippocampal tissues.

### Materials and Methods 1

#### Animals

Young adult male Wistar rats (12 weeks old) were purchased from Tokyo Experimental Animals Supply (Japan). All animals were maintained under a 12 h light/12 dark exposure and free access to food and water. The experimental procedure of this research was approved by the Committee for Animal Research of Nihon University. All animals were maintained under a 12 h light/12 h dark exposure and free access to food and water.

#### Chemicals

E2 and T were purchased from Sigma (USA). ^2^H_5_-E2 and ^2^H_3_-T were also purchased from Sigma (USA).

#### Mass-spectrometric assay of steroids

Step 1) Purification of steroids from hippocampal tissues including solid phase columns .

Adult male rat aged 12weeks was deeply anesthetized and decapitated. After removal of the brain, hippocampal tissue was isolated and stored frozen at -80℃ until use. Note that in the present study freshly prepared hippocampal tissue was not used. The preparation of homogenates of frozen-thawed hippocampal tissue in the double distilled water (DDW) was performed using Fingermasher (SARSTEDT, Numbrecht, Germany). Then, MeOH and DDW were added to become 68%(v/v) MeOH. Isotopes (^2^H_5_-E2 and ^2^H_3_-T) were added to homogenates as internal standards. Homogenates were centrifuged at 3000 rpm for 5 min. After removal of precipitates, the supernatant (2.5 mL) was applied to a GL-Pak Carbograph solid phase column (GL sciences, Tokyo, Japan). The column cartridges were pre-washed with 70%(v/v) MeOH. Elution of steroids were performed with 2 ml of 80%(v/v) MeOH in Toluene. After evaporation, the organic extracts were dissolved in Tetraborate pH Standard Solution (pH 9.18) and subjected to an Isolute SLE+ column (Biotage Japan, Tokyo, Japan), and then eluted with 2 mL of Toluene. After drying, the residue was dissolved in 60 μl of MeOH and 40 μl of DDW. The mixture was centrifuged at 3000 rpm. The supernatant was collected and filtered with Millex GV (Merck, Darmstadt, Germany). The filtrate was subjected to LC-MS/MS.

Step 2) Determination of the nT concentration using LC-MS/MS .

The LC-MS/MS system, consisting of a reverse phase LC (Nexera X2 (Shimadzu, Kyoto, Japan)) coupled with an AB Sciex Triple Quad 6500+ system (AB SCIEX、 Tokyo, Japan), was used for T determination. MS/MS system was operated with electrospray ionization in the positive-ion mode. LC chromatographic separation was carried out on a Sunshell C8 column (150 × 4.6 mm, i.d., ChromaNik Technologies Inc., Osaka, Japan). The mobile phase, composed of two solvents, solvent A (0.05vol% acetic acid) and solvent B (acetonitrile), was delivered at flow rate of 0.9 mL/min. Total run time was 11 min. The initial conditions were held at the mixture of solvent A and B (72:28 v/v) for 0.5 min. After injection of 0.010 ml sample, this was followed by a linear gradient to the mixture of solvent A and B (65:35 v/v) for 3.5 min, and to the mixture of solvent A and B (60:40 v/v) for 4.5 min, and to the mixture of solvent A and B (45:55 v/v) for 3 min and then these conditions were maintained for 2 min.

The ionization conditions of LC separated steroids were as follows: ion spray voltage, 5 kV; turbo gas temperature, 600 °C; ion source gas 1 (nebulizer gas), 65 psi; ion source gas 2 (turbo gas), 65 psi; declustering potential, 80V (for T) and 80V (for ^2^H_3_-T). Nitrogen was used as the collision gas in the Q2 collision cell. In the multiple reaction monitoring mode, the instrument monitored the m/z transition, from 289.1 to 97.0 for T, and from 292.1 to 96.8 for ^2^H_3_-T, respectively.

Step 3) Determination of the nE2 concentration using LC-MS/MS .

The same MS/MS system (AB Sciex Triple Quad 6500+ system) was also used for E2 determination. LC chromatographic separation was carried out on a Kinetex Biphenyl 100A column (150 × 4.6 mm, i.d., Phenomenex, CA, USA). The mobile phase, composed of two solvents, solvent A (Water) and solvent B (methanol), was delivered at flow rate of 0.85 mL/min. Total run time was 11 min. The initial conditions were held at the mixture of solvent A and B (45:55 v/v) for 0.5 min. After injection of 0.015 ml sample, this was followed by a linear gradient to the mixture of solvent A and B (22:78 v/v) for 7.5 min, and to the mixture of solvent A and B (1:99 v/v) for 3 min and then these conditions were maintained for 2 min. This system was returned to the initial proportion of the mixture of solvent A and B (45:55 v/v) within 0.01 min and maintained for the final 2 min of each run.

The ionization conditions of LC separated steroids were as follows: ion spray voltage, 5 kV; turbo gas temperature, 600 °C; ion source gas 1 (nebulizer gas), 65 psi; ion source gas 2 (turbo gas), 65 psi; declustering potential, 90V (for E2), and 95V (for ^2^H_5_-E2) . Nitrogen was used as the collision gas in the Q2 collision cell. In the multiple reaction monitoring mode, the instrument monitored the m/z transition, from 271.1 to 145.0 for E2 and from 276.1 to 147.0 for ^2^H_5_-E2, respectively. The limits of quantification for steroids were measured with blank samples, prepared alongside hippocampal samples through the whole extraction, fractionation and purification procedures. The limits of quantification for 17β-E2 and T were 0.3 pg and 1 pg of hippocampal tissue, respectively.

From the calibration curve using standard steroids dissolved in blank samples, the linearity was observed between 0.1 pg and 1000 pg for E2, and between 0.5 pg and 1000 pg for T, respectively.

#### Statistical analysis

Statistical analysis was performed. Data expressed as mean ± standard error. The number of independent experiments from different animals (n) was used.

### Results 1

The concentrations of nT and nE2 in hippocampal tissues are observed as follows.

<u>nT concentration (average ± SEM)</u>

2.23 ± 0.04 (ng /g wet tissue), 7.8 ± 1.2 (nM) (n=6)

<u>nE2 concentration (average ± SEM)</u>

15 ± 8 (pg /g wet tissue), 55 ± 29 (pM) (n=5)

Using the average hippocampal volume of 0.124 mL (deduced from 0.124± 0.002 wet weight for one whole hippocampus of 12 weeks old male rat, n = 44), the molar concentrations of E2 and T in the hippocampus were calculated. Based on these considerations, 15 pg/g wet weight of E2 in the hippocampus corresponds to 55 pM. The observed nE2 concentration is very different from those obtained in freshly prepared male hippocampus. For example in freshly prepared male hippocampus, nE2 concentration was 2.3 (ng/g wet tissue) [i.e., 8.4 (nM)], and nT concentration was 4.9 (ng/g wet tissue) [i.e., 16.9 (nM)] (Hojo et al 2009). Although a considerable difference (nearly 150 fold) was observed in nE2 concentration between fresh and frozen-thawed hippocampus, nT concentration did not show a considerable difference between these two preparations. These observations indicate that the correct concentration of nE2 is probably very difficult to determine using frozen-thawed hippocampus.

### Discussion 1

Many laboratories have been using “frozen-thawed” hippocampal tissues as starting materials for extraction of steroids, except for Kawato’s laboratory and collaborating laboratories (Hojo et al 2004) (Hojo et al 2009) (Munetsuna et al 2009) (Kato et al 2013) (Yamazaki et al 2013) (Caruso et al 2013) (Li & Gibbs 2019) (Jalabert et al 2022).

The freeze-thaw processes using deep freezer (-80 degree) may damage brain tissues and cells due to crystallization of intracellular water (in freezing processes) and melting of intercellular ice (in thawing processes). Crystallization of intracellular water by freezing may break cell membranes, leading to leaking and losing of antioxidants (such as glutathione and vitamin C) and antioxidant enzymes (such as superoxide dismutase and glutathione peroxidase). Reactive oxygen species produced from mitochondria may easily attack lipids in cell membranes in the absence of these antioxidants and antioxidant enzymes. Melting of intercellular ices may also break cell membranes, leading to leaking and losing of antioxidants and antioxidant enzymes. Under these bad conditions, membrane lipid peroxidation would occur, leading to oxidation of OH at C-3 position of nE2, resulting in inactivation of nE2 molecules. Therefore the risk of oxidation and degradation of nE2 may increase in frozen-thawed hippocampal tissues.

It should be noted that, however, nT has an oxygen (O) at C-3 position which is resistant to oxidation and inactivation. Therefore, nT molecules might not be degraded by freeze-thaw processes of hippocampal tissues. In fact, using freshly prepared (but not frozen-thawed) hippocampal tissues, we always observed the nE2 concentration higher than 1 nM (Hojo et al 2009), (Munetsuna et al 2009) (Yamazaki et al 2013) (Kato et al 2013).

In order to test “freeze-thaw-inducing E2 oxidation hypothesis”, we here measured nE2 and nT using frozen-thawed rat hippocampus. We obtained considerably low nE2 concentration approxi. 15 pg/g tissue (∼55 pM) which is extremely lower than 2.3 ng/g tissue (8.4 nM) in fresh hippocampal tissue (Hojo et al 2009) (Kato et al 2013) (Hojo & Kawato 2018). The nT concentration of 2.2 ng/g tissue (7.8 nM) in frozen-thawed male rat hippocampus was not considerably different from that observed in fresh male hippocampus (∼4.9 ng/g wet tissue, ∼16.9 nM) (Hojo et al 2009) (Hojo & Kawato 2018) .

As another method, a significant nE2 production capacity of P450arom in the hippocampus may be measured by using hippocampal microsomes prepared from frozen-thawed brain tissues. Fortunately, P450aromatase (P450arom) in hippocampal microsomes may not be seriously inactivated by freeze-thaw processes of whole hippocampal tissues. Relatively high P450arom activity was observed in microsomes which were prepared from frozen-thawed female rat hippocampus (Li et al 2016). By using these microsomes, the E2 production activity of P450arom was measured with the application of exogenous T as a substrate. The activity of rat hippocampal microsomal P450arom was approximately 1/10^th^ of the activity of ovarian microsomal P450arom (Li et al 2016). These results support the possibility of high nE2 production in adult rat hippocampus.

Interestingly, in many publications, when the applied concentration of E2 was much lower than 1 nM (e.g., 0.1 nM), E2 did not show rapid nongenomic modulation of LTP or spinogenesis in hippocampal slices from adult rats (e.g., 3 month old) (Warner & Gustafsson 2006) (Mukai et al 2007) (Hasegawa et al 2015). For example, E2 with picomolar concentration (< 100 pM) was not able to rapidly modulate spinogenesis and LTP (Mukai et al 2007) (Hasegawa et al 2015). Note that in hippocampal slices from very young rats (e.g., 1 ∼ 5 week old), 100 pM could be effective to increase Excitatory Postsynaptic Current (EPSC) (Kramár et al 2009).

Achievements of high sensitivity in determination of nE2 is very essential. Several methodological improvements have been performed previously. Examples of methodological improvements were as follows: (A) Organic solvent extraction using hexane, ethyl acetate or methanol in order to have sufficient extraction of neurosteroids from brain tissues. The fresh brain tissue should be used for efficient steroid extraction, and frozen-thawed brain tissues should ve avoided. (B) Pre-purification and separation of neurosteroids with high-performance liquid chromatography (HPLC) (either normal phase HPLC or reverse phase LC) is often used to separate steroids of interest from contaminating fats and other steroids. Solid-phase column is also useful to separate steroids of interest from contaminating fats and other steroids, but efficiency may not be sufficient compared to normal phase HPLC. (C1) LC-MS/MS determination improves sensitivity in determination of target steroids. (C2) Derivatization of E2 at C-3 position, for example, before application to LC-MS/MS could improve the sensitivity more than 10-fold. Specific derivatization of E2 at C-3 position with penta-fluorobenzene (Hojo et al 2009) or dimethylimidazole-sulfonyl chloride (DMIS) (Jalabert et al 2022) or other derivatives (Ebner et al 2006) (Higashi et al 2006) (Li & Gibbs 2019) was very effective. Additional derivatization of steroids at C-17 position with picolinic acid could further improve sensitivity of E2 and T determination, because of induced ionization of picolinic part (Yamashita et al 2007a, Yamashita et al 2007b).

Concerning mouse nE2, another difficulty exists in determination of nE2 concentration. Because the volume of mouse hippocampus is only ∼1/10 of that of rat hippocampus, the number of nE2 molecules are much lower in single mouse hippocampus than those in single rat hippocampus. Therefore, the accuracy of determination of mouse nE2 concentration is much lower compared to rat nE2 concentration. More than four adult mouse hippocampal tissue may be necessary in order to have sufficient accuracy for nE2 determination.

## Section 2: Rethinking dendritic spines in castrated hippocampus

### Introduction 2

Modulation of dendritic spines by androgen in the hippocampus is an attractive theme and has been extensively investigated in castrated male rats (Leranth et al 2003) (Kuwahara et al 2021). In male, castration is used to remove the effects of circulating T and DHT by removing testicles.

It was found that castration induced a significant loss of hippocampal spine-synapse in CA1 region, although no significant effect on the number of pyramidal cells (Leranth et al 2003). Here spine-synapse is the spine which forms synaptic contact as observed by electron microscopic analysis. The density of CA1 spine-synapses was increased (recovered) upon subcutaneous (s.c.) injection of DHT and T (1.6 mg /kg body, 500 μg/day) for 3 days in castrated male rats in which the spine-synapse density was already decreased by castration (Leranth et al 2003).

Injection of flutamide (a blocker of androgen receptor (AR)) suppressed these DHT and T effects in castrated male rats (MacLusky et al 2004). Injection of E2 (15 μg E2/kg body, 10 μg/day) did not show the recovery of the low level of spine-synapse in castrated male to the normal level in intact male (Leranth et al 2003). This could be explained that low E2 dose cannot recover the castrated-rat spine level back to the intact-rat spine level.

The effects of T on spatial memory *in vivo* using castrated male rats were also investigated with behavioral test in combination with Testosterone Propionate (TP)-supplementation (Wagner et al 2018). In order to improve spatial memory, castrated male rats were treated with TP by s.c. injection for 7 days, and were undergone a radial-arm maze test. Dose response of TP supplementation was not simple. T-induced improvement was observed at T dose with 0.125 mg and 0.5mg, but not 0.25mg and 1.0 mg.

Importantly, castration of male rats can considerably decrease nT concentration by 20-30% in rat hippocampus, but surprisingly castration cannot significantly decrease nE2 concentration (Hojo et al 2009), when castration was performed by expert of animal surgery. Therefore, castration-induced deprivation of circulating T alone may not be able to significantly impair the local nE2 action, since male rat hippocampus has both androgen and estrogen signaling pathways that maintain the spine density (Hasegawa et al 2015) (Hatanaka et al 2015) (Koss & Frick 2019). Therefore, T supplementation alone might not be able to sufficiently improve spatial memory performance of castrated male rats.

Not only in isolated slices but also *in vivo*, nE2 plays essential role in modulation of neuronal plasticity and cognitive performance. Rapid and non-genomic modulation by nE2 occurs in addition to genomic modulation. In both male and female hippocampal neurons, nE2 modulates spinogenesis. It was found that the rapid E2-induced increase (within 30 min) of the CA1 spine-synapse density occurred in OVX female rats upon s.c. injection of relatively high dose E2 (45 μg E2/kg body) (MacLusky et al 2005) (Kuwahara et al 2021). However, such an increase in the spine-synapse density did not occur upon injection of low dose E2 (15 μg E2/kg body).

In Section 2, we re-analyzed the castration-induced changes in the density of spines.

Surprisingly, we found that the degree of spine decrease by castration was dependent strongly on skills of castration operators.

### Materials and Methods 2

#### Animals

Young adult male Wistar rats (12 week old, 320-360 g) were purchased from the Institute for Animal Reproduction (Ibaraki, Japan). Castration and sham operations were performed two-weeks before (at 10w old) the experiments of dendritic spine imaging. Castration operation was performed by either the expert of animal surgery (with nearly 20 years of experience) or by Ph.D. students (with about half a year of experience). All animals were maintained under a 12 h light/12 dark cycle and free access to food and water. The experimental procedure of this research was approved by the Committee for Animal Research of Teikyo University.

#### Hippocampal slice preparation (perfusion-fixed slices)

Hippocampal slices were prepared from a 12 week-old male rat deeply anesthetized and perfused transcardially with PBS (0.1 M phosphate buffer and 0.14 M NaCl, pH 7.3), followed by fixative solution of 3.5 % paraformaldehyde. Decapitation was performed at 10:00 – 10:30 a.m. Immediately after decapitation, the brain was removed from the skull and post-fixed with fixative solution. Hippocampal slices, 400 μm thick, were prepared with a vibratome (Dosaka, Japan).

#### Current injection of neurons by Lucifer Yellow

Neurons within slices were visualized by an injection of Lucifer Yellow under a Nikon E600FN microscope (Japan) equipped with a C2400–79H infrared camera (Hamamatsu Photonics, Japan) and with a 40× water immersion lens (Nikon). A glass electrode was filled with 5% Lucifer Yellow, which was then injected for 15 min using Axopatch 200B (Axon Instruments, USA). Approximately five neurons within a depth of 20 - 30 µm from the surface of a slice were injected with Lucifer Yellow (Duan et al 2002).

#### Confocal laser microscopic imaging and analysis

The imaging was performed from sequential z-series scans with super-resolution confocal microscope (Zeiss LSM880; Carl Zeiss, Germany) using Airy Scan Mode, at high zoom (× 3.0) with a 63 × oil immersion lens, NA 1.45. For Lucifer Yellow, the excitation and emission wavelengths were 458 nm and 515 nm, respectively. For analysis of spines, three-dimensional image was reconstructed from approximately 30 sequential z-series sections of every 0.45 µm. The applied zoom factor (× 3.0) yielded 23 pixels per 1 µm. The confocal lateral resolution was approximately 0.14 µm. The z-axis resolution was approximately 0.33 µm. Our resolution limits were regarded to be sufficient to allow the determination of the head diameter of spines in addition to the density of spines. Confocal images were deconvoluted with the measured point spread function using Processing Mode of LSM880.

The density of spines as well as the head diameter were analyzed with Spiso-3D (automated software calculating mathematically geometrical parameters of spines) developed by Bioinformatics Project of Kawato’s group (Mukai et al 2011). Spiso-3D has an equivalent capacity with Neurolucida (MicroBrightField, USA), furthermore, Spiso-3D considerably reduces human errors and experimenter labor. The single apical dendrite was analyzed separately. The spine density was calculated from the number of spines along secondary dendrites having a total length of 40-60 µm. These dendrites were present within the stratum radiatum, between 100-200 µm from the soma. Spine shapes were classified into three categories as follows. (1) A small-head spine, whose head diameter is smaller than 0.4 µm. (2) A middle-head spine, which has 0.4-0.5 µm spine head. (3) A large-head spine, whose head diameter is larger than 0.5 µm. These three categories were useful to distinguish different responses upon kinase inhibitor application. Small-, middle-, and large-head spines probably have different number of α-amino-3-hydroxy-5-methyl-4-isoxazolepropionic acid (AMPA) receptors, and therefore these three types of spines might have different efficiency in memory storage. The number of AMPA receptors (including GluR1 subunits) in the spine increases as the size of postsynapse increases, whereas the number of N-methyl-D-aspartate (NMDA) receptors (including NR2B subunits) might be relatively constant (Shinohara et al 2008). Because the majority of spines (> 93∼95 %) had a distinct head, and stubby spines and filopodia did not contribute much to overall changes, we analyzed spines having a distinct head.

#### Statistical Analysis

Dendrite images were used for spine analysis, and typical images were shown in Fig 1A. Each dendrite has approx. 50 μm in length including approx. 50 spines. For statistical analysis in Fig.1B the Tukey-Kramer multiple comparison’s test was used. We used approximately 50 dendrites with 2300-2700 spines obtained from 3 rats, 12 slices, 30 neurons.

**Figure 1.**
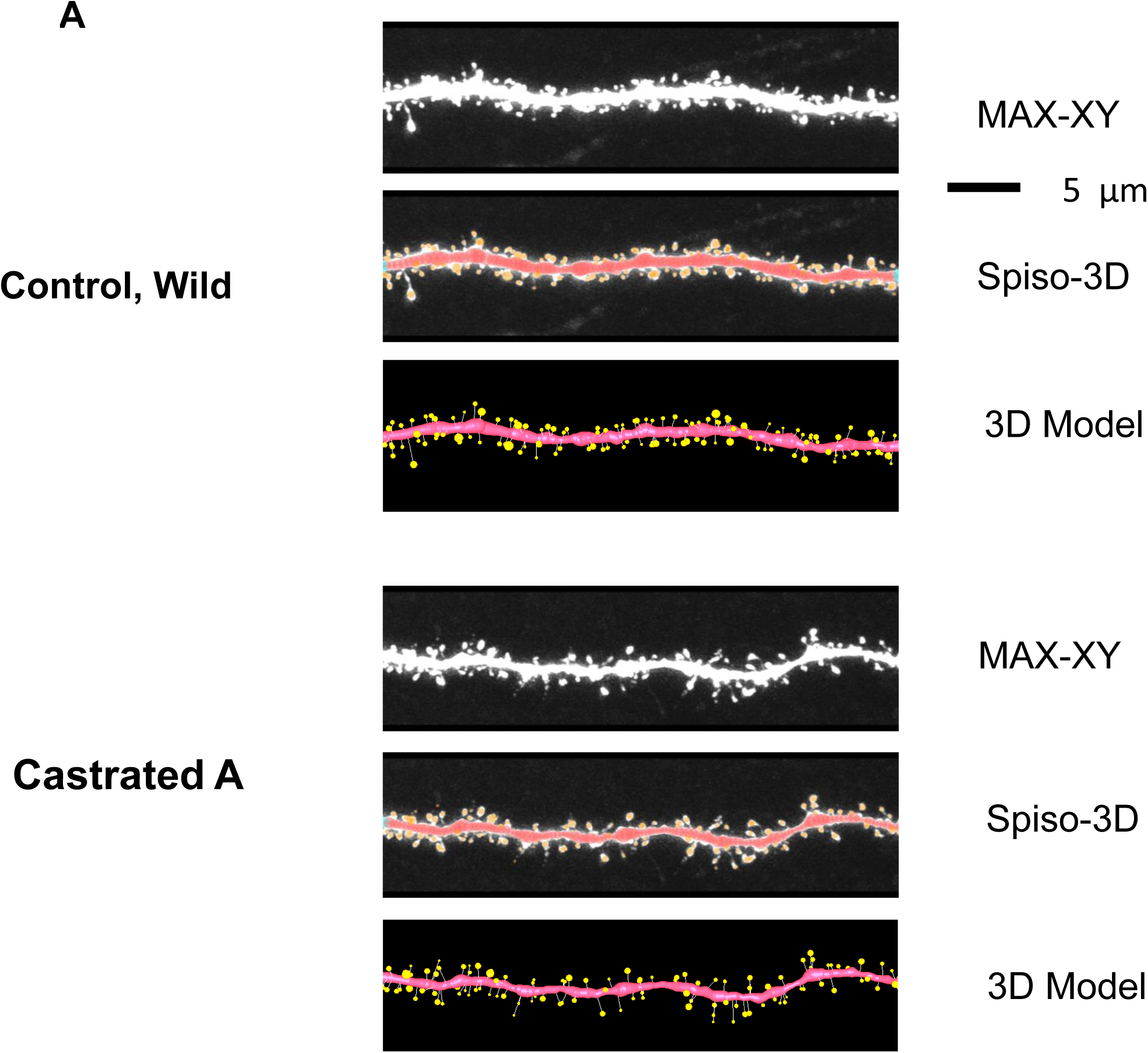
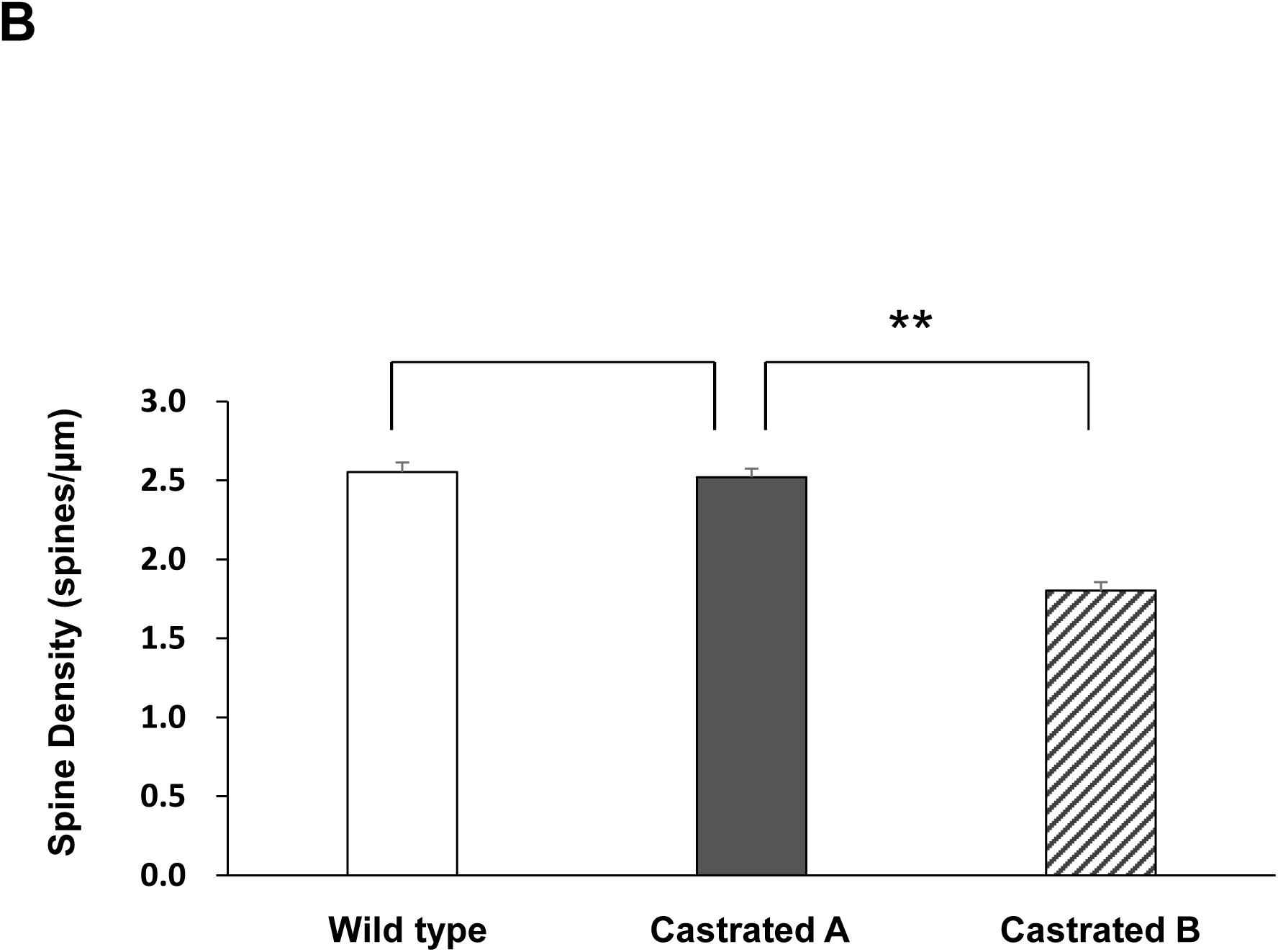
Confocal image analysis of dendritic spines along the secondary dendrites in adult hippocampal CA1neurons. Spines were analyzed along the secondary dendrites of pyramidal neurons in the stratum radiatum of CA1 neurons. Dendrite without treatments (Wild type, Control), dendrite after castration by an expert operator (Castrated A type), dendrite after castration by PhD students (Castrated B type). **(1A)** Upper image (MAX-XY) shows maximal intensity projections onto XY plane from z-series confocal micrographs. Middle image (Spiso-3D) shows the dendrite and spines analyzed with Spiso-3D software. Traced dendrite is shown with connected series of red balls and spines are indicated with yellow balls. Lower image (3D Model) shows 3 dimensional model illustration of (Spiso-3D) image. Bar, 5 μm. **(1B)** Comparison of the total spine density between Wild type, Castrated A type and Castrated B type. Vertical axis is the average number of spines per 1 μm of dendrite.

### Results 2

We measured the effects of depletion of circulating androgen on dendritic spine density by castration of adult male rats.

We used two types of castrated rats, one is Castrated A type and the other is Castrated B type (Fig. 1). Castrated A type was prepared by an expert operator having nearly 20 years of experience of castration surgery. Castrated B type was prepared PhD students having nearly half a year experience of castration surgery.

As shown in Fig. 1B, CA1 total spine density was not different at all between Castrated A and Wild type control. In contrast, spine density was significantly decreased in Castrated B. One possible reason is that because of relatively poor skills, PhD students may inhibit blood circulation for relatively a long time (> 5 min) during removal of testis, resulting in a micro brain ischemia leading to decrease in spines but not inducing neural death. However, an expert technician did not inhibit blood circulation for more than 1 min, therefore a micro brain ischemia does not occur.

From the steroid concentration analysis of Castrated A type rats (Hojo et al 2009), the hippocampal T level was significantly decreased to approxi. 20 – 30%, and the DHT level became 0% in Castrated A rats. In blood plasma, T and DHT were completely depleted in Castration A.

Fig. 2 shows that the morphological changes in the spine head diameter distribution were not observed between Castrated A and Wild type control. No changes were observed in the relative population of small head spines, middle head spines and large head spines.

**Figure 2.**
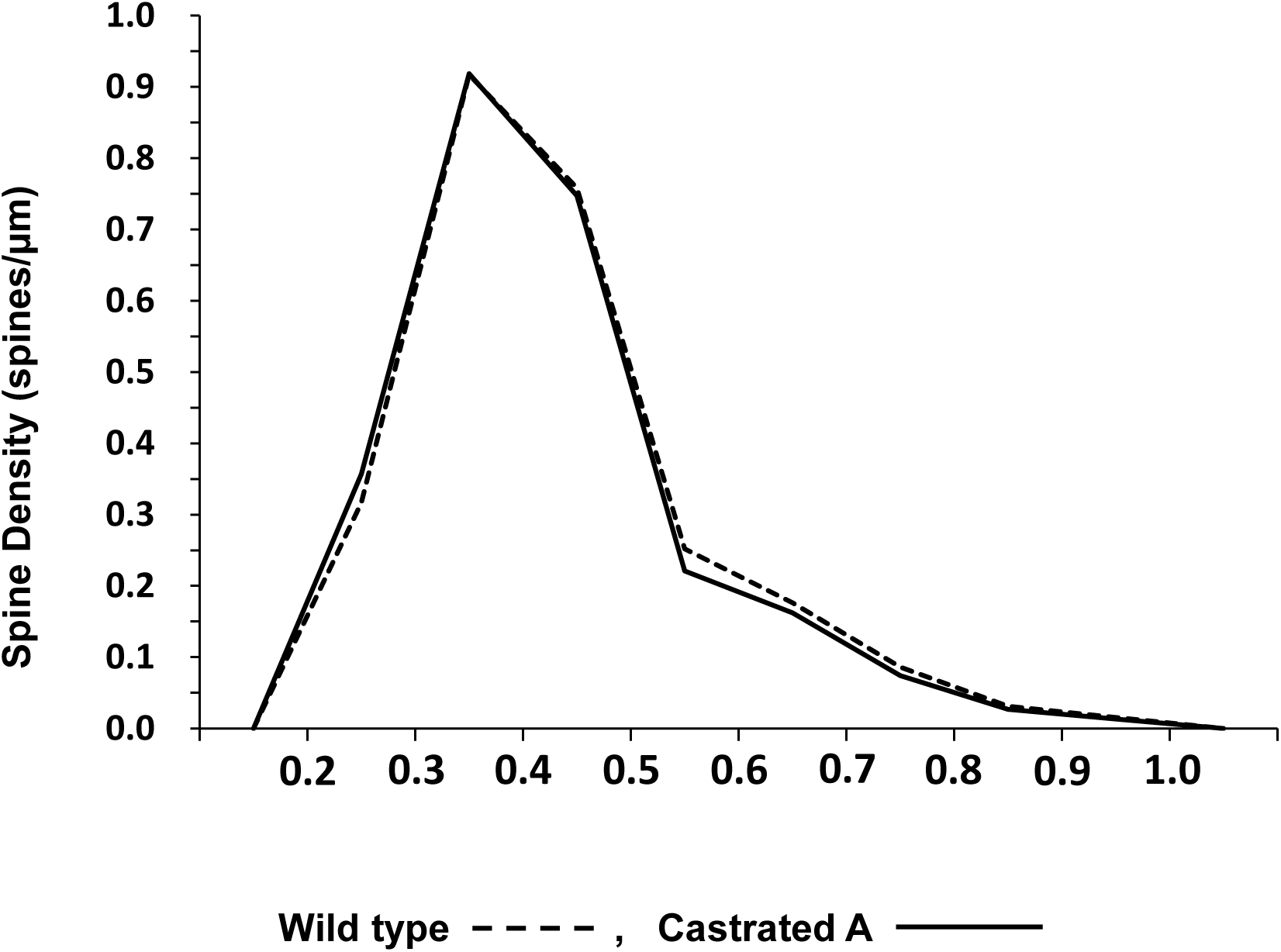
Morphological analysis for effects of castration on spine head diameters in adult hippocampal CA1 neurons. Comparison of spine head diameter distribution between Wild type (control, black line) and Castrated A type (dashed line). Vertical axis is spine density (spines/μm), and horizontal axis is spine head diameter (nm).

### Discussion 2

In Castrated A type hippocampus, a considerable decrease in nT concentration (to 20-30% level) was observed, but no significant decrease in nE2 was shown (Hojo et al 2009). Therefore, castration-induced deprivation of circulating T alone cannot considerably impair hippocampus-dependent behavior supported by local nE2 action. Since male rat hippocampus has both androgen and estrogen signaling pathways that modulate the spine density and synaptic plasticity, not only blocking AR (androgen receptor)-signaling but also blocking of ER (estrogen receptor)-signaling are necessary to completely block sex-hormone effects (Hasegawa et al 2015) (Hatanaka et al 2015) (Koss & Frick 2019). Therefore, T supplementation alone might not be able to considerably recover the spatial memory in castrated male rats.

## Abbreviations

nE2: neuro-estradiol
nT: neuro-testosterone
LC-MS/MS: liquid chromatography with tandem-mass-spectrometry

## Acknowledgements

We thank Mr. Hiroyuki Ochiai and Mr. Tetsuya Imazu (ASUKA Pharmaceutical Co. Ltd.) for steroid concentration determination.

## Author Contributions

SK conceived and designed the study. MO-I, MS and KM conducted the experiments and analysis of the data. SK wrote the manuscript. SH and MS contributed to discussions. All authors provided feedback on the manuscript.

## Conflict of Interest Statement

The authors declare no conflict of interest.

## Notes

### Competing Interest Statement

The authors have declared no competing interest.

### Summary of Updates

Figure 1A is revised by adding Castrated A figure.

